# HandMol: Coupling WebXR, AI and HCI technologies for Immersive, Natural, Collaborative and Inclusive Molecular Modeling

**DOI:** 10.1101/2023.11.24.568613

**Authors:** Fabio J. Cortes Rodriguez, Lucien F. Krapp, Matteo Dal Peraro, Luciano A. Abriata

**Author notes:** Corresponding autor.

## Abstract

Except for isolated developments and specific software extensions, molecular graphics and modeling have historically been stuck at flat screens for visualization, mouse operations for molecular manipulation, menus and command line interfaces for controls, and single-user interfaces that only allow collaboration by streaming video hence limited to just sharing the view of the user operating the software. We demonstrate here how various technologies are ripe enough to enable much more fluent, immersive and natural human-computer interactions that in turn facilitate collaboration between human users, using affordable hardware through the internet and without even installing any specialized programs. For this, we introduce HandMol, a web app that exploits (i) WebXR for molecular visualization and manipulation in virtual reality, (ii) speech recognition coupled to a large language model to pass commands orally, (iii) speech synthesis for auditory feedback, (iv) WebRTC to communicate multiple instances of the tool without even requiring a server, and (v) external APIs to flexibly account for molecular mechanics, exemplified here with an endpoint running an AMBER forcefield for protein and nucleic acids and another running a DFT-trained neural network, ANI-2x, to allow exploration of conformation and some simple reactivity at high speed and accuracy. We show example applications to situations from daily work and education in chemistry and structural biology where HandMol can provide an advantage over traditional software: exploring and explaining molecular conformations and reactivity, docking and undocking small molecules into/out of protein pockets, threading molecules through nanopores, preparing systems for molecular simulations and for protein design, etc. We also present a brief study showing how users, even with limited or even no experience in VR, can significantly benefit from these kinds of technologies. As a draft prototype for the moment, HandMol is made available free of charge and without registration at https://go.epfl.ch/handmol, in (optional but greatly appreciated) exchange for feedback on usability and on features expected for this kind of tools.

## Introduction

In disciplines like chemistry, materials science and structural biology, human ability to visualize and manipulate molecules in three-dimensional space is of utmost importance. Traditional molecular graphics software solutions relying predominantly in two-dimensional mouse moves and visualization to convey spatial information of intrinsically 3D nature, have long served as the foundation for computational chemistry, molecular modeling, and structural biology. However, these setups are very limited in their capacity to offer depth perception and spatial manipulation, especially regarding operations aimed at editing atom coordinates such as displacing molecules, breaking or forming bonds, changing conformations, or simply pointing at specific features of a structure. Additionally, manipulation of molecular structures in traditional software is also limited by the user’s need to master commands and keyboard/mouse shortcuts. Besides, traditional interfaces limit operations to single users acting with single hands, which are suboptimal for engaging discussions and for finely manipulating atom coordinates. In particular, discussion on molecular structures is limited in that a single operator can control the views, and other people remain as passive observers.

In recent years, four technologies emerged that could be applied directly to address the limitations of traditional software for molecular graphics and modeling. First, powerful hardware for virtual reality (VR) and augmented reality (AR) that is also more affordable and wearable and counts with environment and body tracking capabilities plus Wi-Fi connectivity, now enables robust, deeply immersive experiences and two-handed interaction with objects via the user’s bare hands or via handheld controllers. Second, despite not widely used, speech recognition has matured enough to be of actual help in simplifying human-computer interactions, as exemplified in an application to voice-controlled quantum chemistry based on modular integration.^1^ Third, large language models further facilitate (besides enabling many other applications) voice control by casting requests in natural language to commands and scripts, as exemplified in a demo showing how to control the JSmol plugin for molecular visualization via spoken commands, assisted by GPT-3.^2^ Fourth, machine learning models that can parse molecules and compute energies and forces allow today fast, simple and accurate estimation of molecular mechanics, that could help to inform interactive software for molecular graphics and modeling seamlessly. For example, the ANI-2x neural network can compute DFT-quality energies and forces orders of magnitude faster than possible with actual DFT calculations.^3^

Critical to the work presented here is that all these modern technologies exist in forms that can be easily integrated together in a modular fashion. This enables the straightforward development of software with complex features, particularly for the web. Here we present a first integration of the above-mentioned technologies into a draft prototype of an app that runs in the web browsers of VR headsets, smartphones, tablets and computers, allowing one or two users to visualize and manipulate molecules in immersive 3D while interacting with the hardware and software as naturally as possible with current technologies (Figure 1). We have called this web app HandMol, and it is available out of the box on all devices by pointing their web browsers to https://go.epfl.ch/handmol.

**Figure 1.**
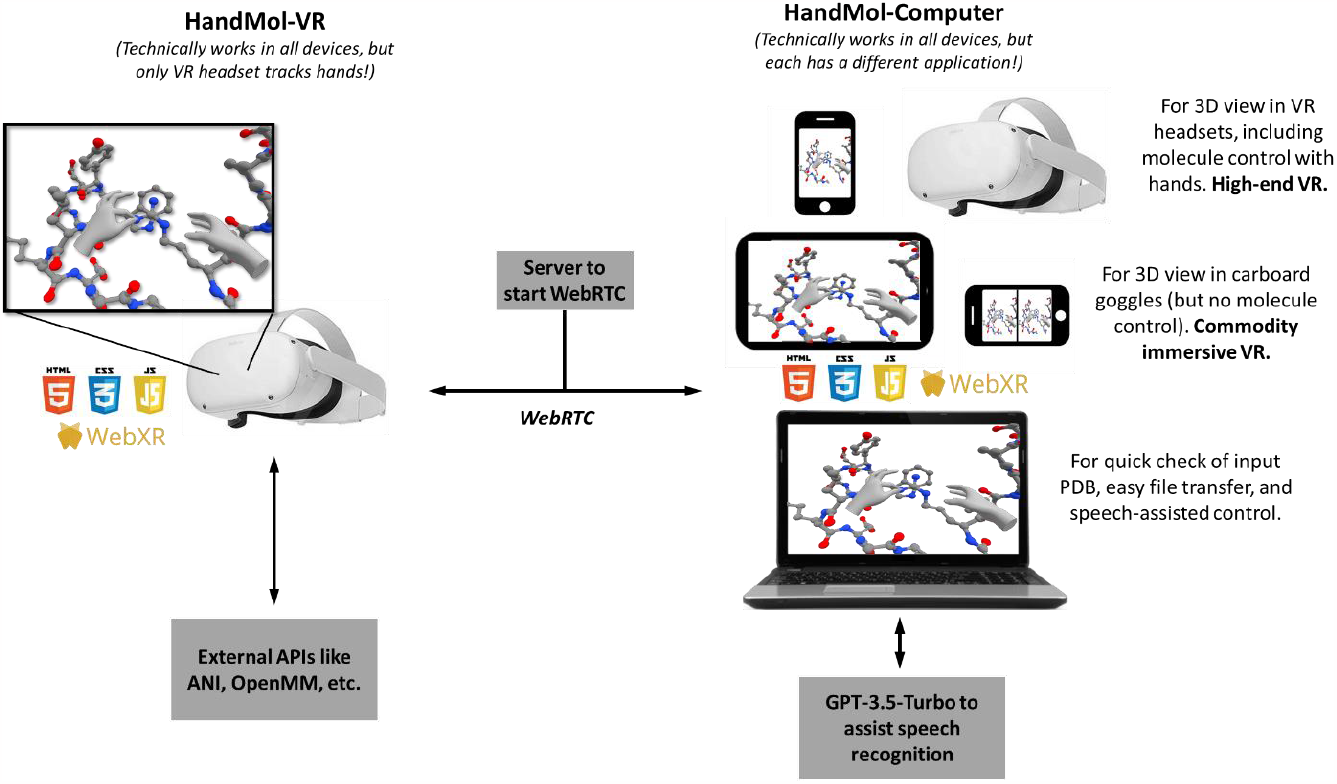
HandMol architecture. Scheme showing how HandMol’s components, called HandMol-VR and HandMol-Computer, are built from HTML, JS, CSS, WebXR, WebRTC, and API calling components; and how they can be used in different scenarios and with different pieces of hardware. The view is a true screenshot from inside a VR headset, where the background was removed and painted white for simpler visualization in this schematic figure. For the untouched screenshot, check Figure 2A; also check other figures of this article and try out the examples at https://go.epfl.ch/handmol.

### Virtual reality for molecular graphics and modeling

Over the various waves of “hype” on AR and VR, these technologies attempted a few times to irrupt in the field of molecular graphics and modeling, with very limited success and even some reports of no actual benefit in research.^4^ Although these negative results and lack of impact can be debated and certainly depend on many details whose analysis exceeds the scope of this work, there is no doubt that in practice, AR and VR solutions had very limited acceptance and penetration into the target communities of researchers and educators. We pose that leaving aside a matter of costs of the VR devices, an important limitation to the adoption of AR/VR software for molecular graphics and modeling arises from the steep learning curves required to use these devices, the fatigue that some people can experience in VR, and certainly the limited features offered by existing software. However, costs are quickly dropping, with several devices now reaching the prices of consumer smartphones. VR fatigue is much less of a problem with modern headsets (for example, in a recent work we had over 150 people aged 12-80 experiencing 15-20 min VR sessions with no problems^5^). Usability has also made large progress in the last 3-5 years, thanks to detailed onboarding, simplified all-in-one hardware with no cables or external computers and cameras required, and simplified controller operation or even no-controller operation for devices that support hand tracking.

We pose that although the state of the art is not yet ready to fully replace computers for the main workload in molecular graphics and modeling, and that this might not even be close to happening or even “make sense”, VR technologies are ripe enough to facilitate the hardest, most “3D-intensive” operations and discussion to be carried out in a new, far more efficient way. This is exactly where HandMol and the technologies discussed here fit.

At the core of its design, HandMol attempts to solve the following problems of current VR software for molecular graphics: how to easily move molecules back and forth between VR headset and computer, how to account for molecular mechanics realistically but inexpensively, how to control commands without a traditional interface at hand, and how to run not in just 1 or 2 devices but in an as wide array of devices as possible. We have tackled these problems by capitalizing on the WebXR standard and API (see below), speech recognition and speech synthesis assisted by OpenAI’s GPT-3.5-Turbo large language model to cast oral commands into functions, WebRTC to wirelessly connect the VR headset to a computer through which the user moves atom coordinates with the alternative use of coupling users inside VR, and some novel ways to simplify molecular mechanics complemented with high-quality feedback from calls to external forcefields.

In the next sections we present our first prototype of HandMol, which is accessible free of charge and without registration at https://go.epfl.ch/handmol in exchange for feedback on usage and suggestion of features expected for such a tool. We first overview how HandMol was built, and we then explain how it works. Next we showcase example applications to situations from research and education in chemistry and structural biology, illustrating the exploration of molecular conformations, docking and undocking small molecules into protein binding pockets, threading molecules through nanopores, knotting molecular structures, setting up systems for molecular simulations, probing simple reactivity, building molecules, etc. We conclude by discussing further extensions of the technology and HandMol itself, via prototypes that run virtual or augmented reality in VR headsets and in regular smartphones. Every single figure shown here can be reproduced right away by accessing HandMol and choosing the right example; and even readers without a VR headset can try HandMol on a laptop, tablet or smartphone.

### System

#### Core components

One particular development that has been rapidly expanding the use of VR and AR in the last years is the emergence of WebXR, a standard and API that facilitates cross-device and cross-OS experiences of augmented, virtual, or mixed nature (AR/VR/MR) directly inside web browsers, spanning from VR headsets to smartphones, tablets and computers. WebXR greatly simplifies programming and maximizes reach, because a single piece of code shall in principle work in all the devices that follow the standard -including all the modern headsets from Meta, Apple, HTC, Pico, Lenovo and other companies. On the user side, the WebXR technology certainly facilitates deployment and use, because the app is readily available on the web and as such can be accessed through an URL like any other web page, thus bypassing the need for software installations.

HandMol leverages the power of WebXR for seamless and intuitive molecular manipulation in VR. The current prototype consists in two web pages, called HandMol-VR and HandMol-Computer. Each app can be used separately, to either load and see a molecular system inside VR or in a computer, the former benefiting from VR’s deeply immersive view and the latter from simple loading and saving of atomic coordinates (in PDB format). Linked together through a peer-to-peer (WebRTC-based) connection directly through the internet hence without cables, the two apps allow the user to load coordinates in the computer, send them to the headset to engage in hand tracking-assisted edition of atomic coordinates aided by forcefields, and then saving the modified atomic coordinates in the computer (Figure 1 and 2A).

**Figure 2.**
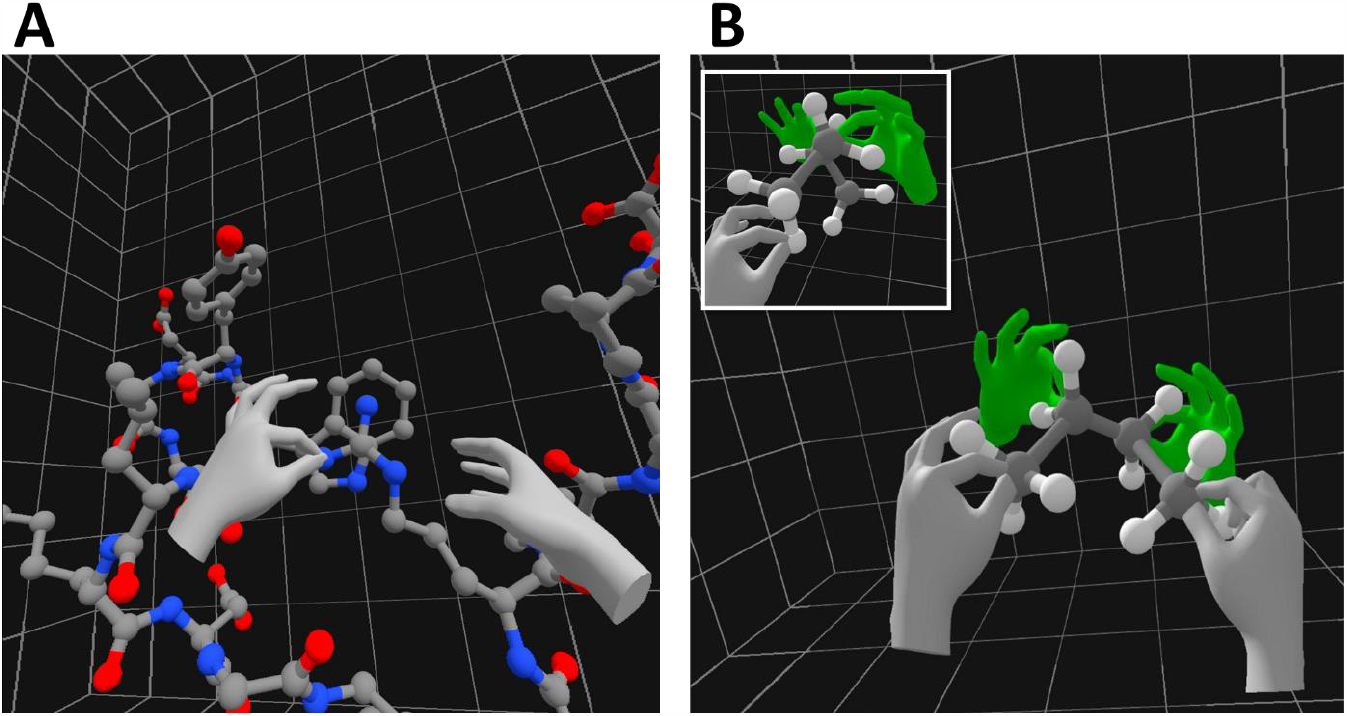
One and two users inside VR with HandMol, and a chance to present more example systems. (A) View of a user running HandMol-VR in a VR headset, here showing how an Arginine sidechain could form a cation-pi interaction with the aromatic rings of a Tryptophan sidechain. (B) View of a user running HandMol-VR in a VR headset and acting with his hands (seen in grey) on a molecular system together with another user who is running HandMol-Computer in a VR headset too (green hands). In this case, the users are together exploring the conformations of butane: while the green user holds two carbons tight, the grey user (whose view is shown) drives rotation around the central C-C bond. The inset shows the a similar operation but seen from a point aligned with the two central C atoms.

Importantly, if the HandMol-Computer app is accessed in a web browser supporting speech recognition and synthesis such as Google Chrome, then all the communication between headset and computer can be done via spoken commands and is assisted by spoken feedback from the computer. For this, the system uses the Web Speech API and calls to OpenAI’s GPT-3.5-Turbo language model to assist speech recognition and casting to commands (this last feature requires that the user provides an OpenAI API key on HandMol-Computer).

An interesting “hack” is to run the HandMol-Computer app in a VR headset too, in which case two users can concurrently view and modify a molecular system. Each user will see his/her own hands, the other user’s hands, and the molecules, on which they can act independently (Figure 2B).

#### Molecular mechanics

To provide visually realistic feedback on molecular mechanics in real time as users move atoms around the space, HandMol employs an approach based on rigid particle physics operated by the Cannon.js library that builds constraints based on molecular structure as in the Virtual Modeling Kits^6^ of our MoleculARweb platform.^7^ For more physically accurate simulations, activated either on call via spoken commands or automatically when the user releases all atoms (if the *Minimize on release* switch button is on), HandMol allows use of either an AMBER forcefield for protein, RNA and DNA molecules, or the ANI-2x DFT-trained neural network^3,8^ for small molecules -both currently carrying out local minimizations only.

ANI-2x minimizations run directly from its PyTorch implementation,^9^ while AMBER minimizations run on OpenMM,^10^ both accessed via a dedicated API we have put in place on a private server. Each minimization is run in 3 repeats of 10 gradient descent steps; when each step is completed HandMol refreshes the visualization and the Cannon engine to convey a graphical depiction of the minimization taking place. The last recalculation of the Cannon restraints (after the third repeat) allows fluid dynamics when handling the molecules in a way consistent with the new conformations generated; besides, this allows covalent modification of molecules (possible with ANI, although not technically devised for this) to remain consistent.

### How to use HandMol -please read this carefully before trying it out

The HandMol version available as of the time of submission of this preprint is only a prototype to demonstrate integration of the technologies, and as such it is fragile and buggy. So please do follow these guidelines to get an optimal experience while trying it out.

#### Single user with VR headset only

For use only in a VR headset without linking to a computer, navigate to https://go.epfl.ch/handmol-vr in the headset’s web browser and choose HandMol-VR. Once the app is loaded, go straight to Step 2. Here, copy-paste a PDB file into the text area labeled *Current PDB* or choose among one of the preset examples, and click *Parse PDB* to see the molecule in the background (this command also sends the PDB to the connected computer if any, but here we are assuming a VR headset-only session).

To get immersed in the molecular system, click *ENTER VR*. Inside VR, the molecule and your hands are visible. By pinching on atoms with the thumb and index fingers (of both hands, even simultaneously) you can move them at will, effectively translating and changing the conformations of the molecules.

Quitting the VR mode brings you back to the main HandMol-VR window, where options can be changed before you go back into VR model. For example, you can activate minimizations with ANI or Amber on atom release, or adapt the scale settings to make the system fit as desired, and then enter back inside VR (Figure 3A).

**Figure 3.**
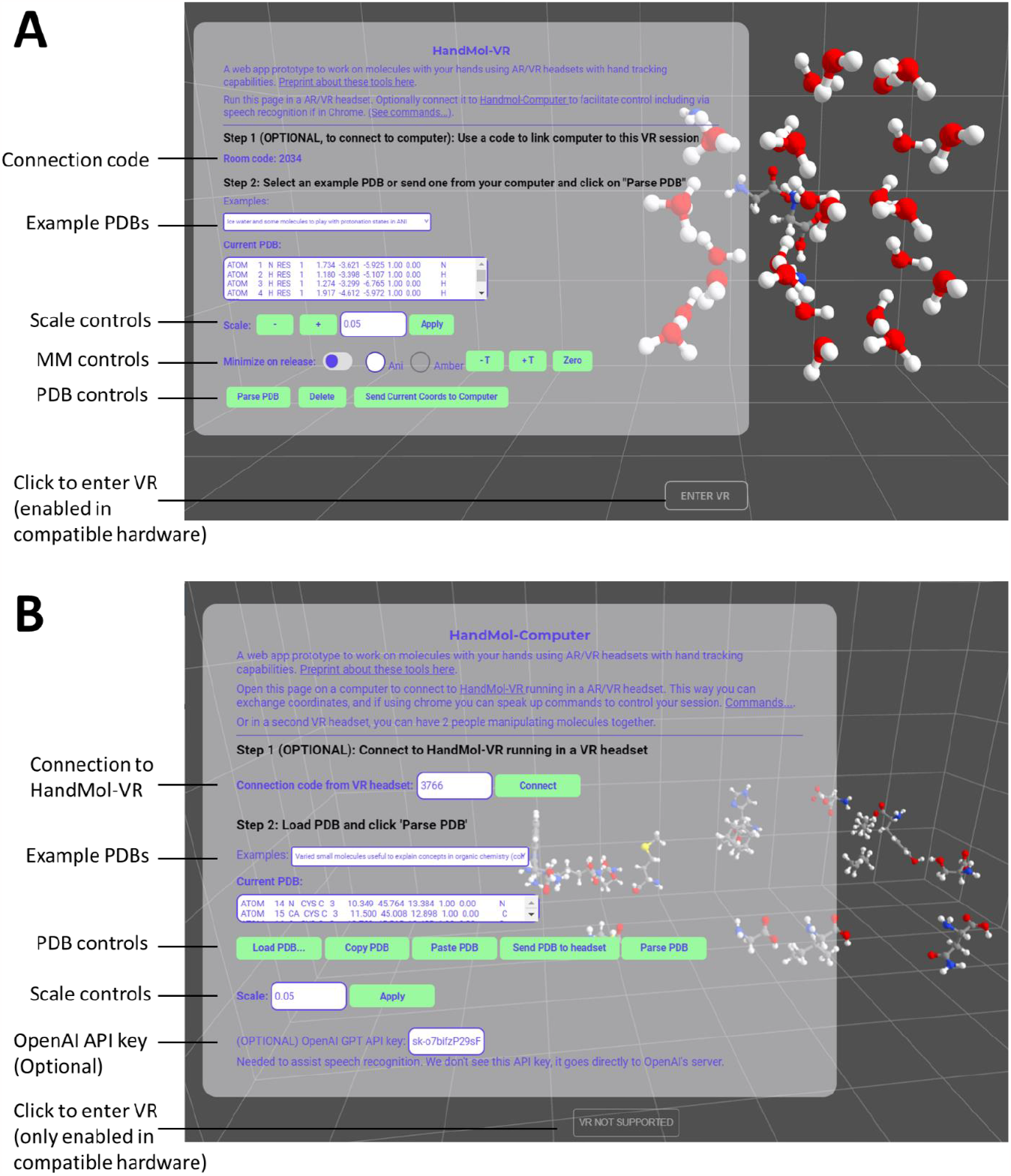
HandMol-VR and HandMol-Computer outside VR mode, with controls exposed. (A) HandMol-VR as seen in the web browser of a VR headset, before accessing VR (i.e. before clicking the *ENTER VR* button at the bottom). (B) HandMol-Computer as seen in the web browser of a laptop computer, which doesn’t allow VR access (hence the *ENTER VR* button displays as *VR NOT SUPPORTED*).

#### Single user with VR headset linked to computer

To use HandMol in a VR headset linked to a computer, open HandMol-VR in the headset’s web browser and HandMol-Computer in a web browser running in a laptop or desktop computer. If you want to benefit from speech recognition and speech synthesis, use Google Chrome in the computer.

With HandMol-Computer you can load or paste a PDB file under *Current PDB* in Step 2, and then do *Parse PDB* to quickly check that it displays correctly (Figure 3B) (this operation also sends the PDB to the VR headset if already connected).

In Step 1 of HandMol-Computer you can enter a 4-digit numerical code that HandMol-VR provides when it loads, and click *Connect* to link your HandMol-VR and HandMol-Computer sessions. Alternatively, you can tell the computer to “*connect with code*” followed by the numerical code (the system also understands variations of this command, especially if an OpenAI API key is provided). Once the connection is established, you can control HandMol-VR by speaking up commands like “*Zoom in*”, “*Relax with Amber*”, “*Send me the coordinates*”, etc. A dedicated menu lists all available commands, which are understood flexibly if a valid OpenAI API key is provided.

#### Two users in VR headsets

In this collaborative mode, two users can see each other’s hands and act with all four hands simultaneously on the system (Figure 2B). To achieve this, one of them must open HandMol-Computer in a VR headset rather than in a table, and connect to the code provided by the other user who is running HandMol-VR in his/her VR headset.

This mode actually works as well if the second user loads HandMol-Computer in a WebXR-enabled tablet or smartphone, the latter further allowing immersive visualization by clicking *ENTER VR* and plugging the smartphone in landscape mode inside cardboard goggles (see under discussion when referring to Figure 5A)

#### Note about PDB files and molecular mechanics in HandMol

While an engine for rigid-body physics provides continuous realistic flexibility in real time, minimization with more accurate means is also at hand. You can activate *Minimize on release* to force minimization with AMBER or ANI when all atoms are released (by both users if two are inside VR). Optionally, you can call for a minimization with a spoken command such as “Minimize with ANI”. For minimizations to run properly, PDB systems must be created with care as explained under “Molecular Mechanics” in the “System” section above, and in what follows.

The PDB coordinates used for input can contain one or more molecules and isolated atoms, as covered throughout the examples shown in this article and available in the app. Approximation of molecular flexibility with the rigid-body physics engine can parse all atoms of the periodic table and is robust to incomplete molecular structures, as in the example of the aerolysin nanopore through which the user can thread a short segment of single-stranded DNA (Figure 4A). An interesting “hack” when using only the rigid-body physics engine is that flagging the Occupancy column with xxxl makes the atom rigid, restraining it to its starting position. This is used in the example allowing to thread atoms through a carbon nanotube, where fixing a bunch of C atoms of the nanotube forces it to stay in place while the user threads the molecule (Figure 4B).

**Figure 4.**
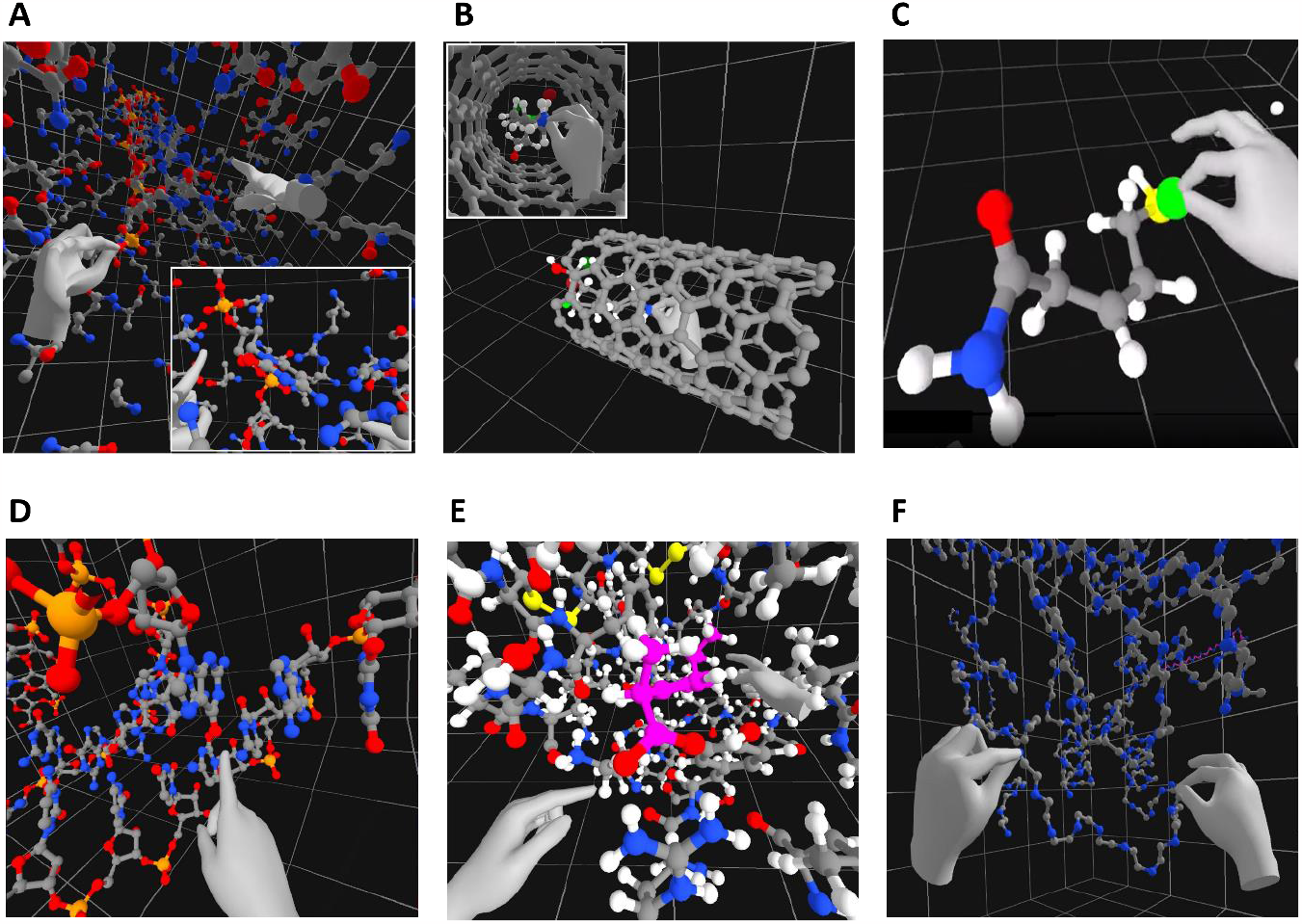
Molecular mechanics in HandMol. (A) A user threading a piece of single-stranded DNA through an aerolysin nanopore assisted by the rigid-body physics engine only, seen from the side where the DNA molecule is headed. The user’s left hand is seen pulling the DNA from a phosphate group, and the right hand is pointing at an Arginine that poised to interact with a DNA base at the reading site. Inset: View closer to the base being read, where the user’s right hand points at the Arginine’s guanidinium group and the left hand points at a phosphate that would be locked to other Arginine sidechains. (B) A carbon nanotube through which a user is pulling a molecule, assisted by the rigid-body physics engine. (C) Building a molecule inside the “Virtual Modeling Kit” example, here approaching an F atom to add it into the growing molecule, assisted by ANI-2x. See video version at https://twitter.com/labriataphd/status/1726726066633343052. (D) Pulling ibuprofen out of its pocket in BSA, assisted by rigid-body physics and running on a portion of an X-ray structure extracted around the small molecule. In this case the representation includes the H atoms, essential to better grasp how the small molecule fills the volume of the pocket and also helping to see the hydrogen bonds and electrostatic interactions the stabilize the complex. (E) Modeling how a peptide could connect two separated pieces of a protein being designed (the tool serving to approximate the length and pose of the required peptide, that will then be modeled in a program like RoseTTAFold Diffusion or similar). (F) A single-chain RNA molecule folded in space, explored with help of an AMBER forcefield.

Contrary to the robustness of the rigid-body-based simulation, if a forcefield will be used then all molecules in the input PDB must be described accordingly - and no error handling is available in the prototype. Namely, to be processed by ANI-2x, all molecules must contain all H atoms, and only atoms supported by this system can be included (H, C, N, O, F, Cl, and S). Meanwhile, Amber can currently only process standard residues making proteins, DNA and RNA, but no ligands, post-translational modifications or non-natural amino acids. Note that the AMBER system as currently implemented reads proteins without their H atoms, even if provided. For calculations, the H atoms are invisibly added on the fly but they are removed for display. Last, a limitation of the AMBER system as of the date of uploading this article as preprint is that it can only treat systems complete with all atoms (for cleaning and completing protein structures, we recommend the Protein Repair & Analysis Server^11^).

For quick tests, HandMol includes a series of preset examples that allow users to quickly test the examples presented here, including some specifically suited for Amber and ANI-2X, with or without connecting to a computer. One of the examples for ANI is a full Virtual Modeling Kit where the user/s can grab atoms from a collection and build molecules, as shown in Figure 4C (video linked in the caption). HandMol also brings three examples of systems tractable with Amber: the fast-folding tryptophan cage (depicted in Figure 2A), a short piece of single-stranded DNA, and a folded RNA molecule (Figure 4D).

There is in this prototype version an important limitation in the size of the tractable systems, especially on the side of HandMol-VR. In the Oculus Quest 2 VR headset, which we have used throughout most of HandMol’s development, the program currently manages smoothly up to around 700 atoms, and can be pushed up to around 1,200-1,300 atoms at the expense of some lag and possible VR sickness. This upper limit corresponds to a protein of around 120-140 residues represented without H atoms (note that Amber minimizations add H atoms on the fly, so lacking these atoms in the protein structure isn’t a limitation when applying this forcefield). In order to work with bigger molecular systems, a possible workaround (but neither ANI nor Amber work with it, only the rigid-body-based system does) is to extract portions of interest as shown in the preset examples for the aerolysin nanopore with a ssDNA molecule ready to be translocated (Figure 4A) or BSA’s binding site filled with ibuprofen (Figure 4E), or the protein design problem where only the core backbone is provided together with polypeptide strands to be grabbed and placed for modeling (Figure 4F).

### Prospect, extensions, and waiting for your feedback

While we envisage fixing HandMol’s several problems and bugs to improve usage and stability, the current version presented here is only meant to showcase the power of coupling all modern technologies, rather than being a closed, stable solution. In the immediate pipeline, we will also work on extending HandMol’s capabilities along a few critical lines: implementing more forcefields via OpenMM, improving and extending the functionalities for voice control and spoken feedback, and exploring how to improve and further control visualization.

If you have tried out HandMol, please let us know what worked and what didn’t work, as well as what features you would expect from a tool like this, by filling the form at https://docs.google.com/forms/d/e/1FAIpQLSeC9rUEgJ_7eXhXqPE7r8FSg5qDvY_xGy-eTZSGqfBBuOYY4w/viewform?usp=sf_link

We aim next to further explore and build on other capabilities of the WebXR API, including mixed-reality modes and usage in goggle-inserted smartphones. VR view using goggle-inserted smartphones is already supported out of the box (Figure 5A), and mixed reality/augmented reality modes can be achieved with few changes in code (Figure 5B and 5C), as shown in the very draft prototypes available at HandMol’s main page (links saying “Mixed reality mode” under the links for HandMol-VR and HandMol-Computer). Indeed, the augmented reality mode also works out of the box in smartphones and tablets (Figure 5D).

**Figure 5.**
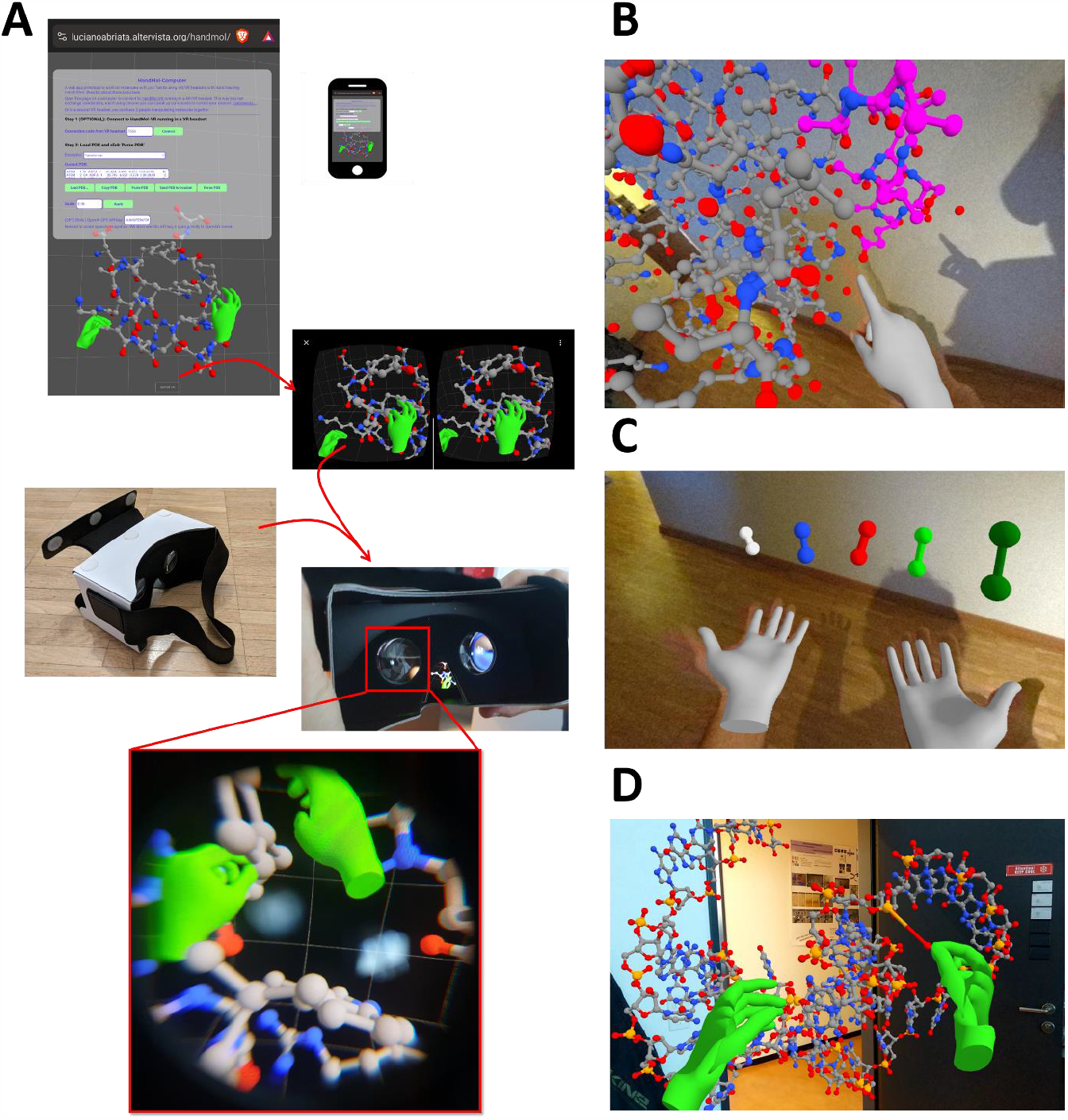
Extended modes: VR in cardboard goggles, high-end mixed reality in modern VR headsets, and commodity augmented reality in smartphones. (A) Accessing either HandMol-VR or HandMol-Computer in a smartphone (best if its browser is set in “Desktop site” to fit all the GUI in the screen) and clicking on *ENTER VR*, results in the WebXR API splitting the view in two for stereoscopy assisted by the device’s motion sensors. Inserting the smartphone inside cardboard goggles immediately allows immersive view much like in a VR headset, with the possibility to inspect the view by freely moving the head (but no displacement is possible, and no hand-tracking is available). (B) Visualizing a protein complex (Ubiquitin bound to an interacting helix, PDB 2D3G) with a VR headset (here Oculus Quest Pro) in mixed reality mode. (C) Building diatomic molecules made of different atoms assisted by ANI-2x, as seen in mixed reality. (D) HandMol-Computer running in Augmented Reality mode on a smartphone linked to another user who is running HandMol-VR in a headset, whose hands are seen in green.

## Discussion

We have here presented a prototype of HandMol, a piece of entirely web-based, mostly client-side software that integrates WebXR, AI methods, and HCI technologies to put forward a new way to visualize and interact with molecules in an immersive and collaborative way, further assisted by computer calculations of mechanics.

These immersive technologies have the potential to bridge the gap between the abstract representations on 2D screens and the tangible reality of three-dimensional molecular structures. VR and AR offer deep immersivity, enabling users to interact with and manipulate molecules as if they were real-world objects, fostering a deeper understanding of their intricate spatial arrangements. Yet so far, VR/AR software of actual utility has been developed only slowly, with many solutions that address different problems and integrate technologies only partially, and suffering from many problems: steep learning curves, especially with older VR hardware (and this means just 5 year old hardware as of 2023), difficulty in moving files in and out of VR headsets, complicated ways to access commands when in VR, difficulty in handling complex menus and buttons in handheld controllers, sometimes VR sickness, etc. As technologies ripe, and as software creators can address all these problems, we are positive that VR can become of actual use in chemistry and structural biology, for education and research, as we intend to communicate through the several examples applying HandMol.

Of course, HandMol is not the first tool to offer VR visualization and manipulation of molecules, with software like Narupa and Nanome standing out. Even web-based solutions for static protein visualization exist that exploit WebXR, like ProteinVR, and also non-web tools that can simulate peptide dynamics quite realistically, like Peppy. We ourselves also presented recently a tool that allows static visualization of molecules in WebXR.^12^ The problem with these tools is that either they treat molecules as whole (rigid) objects, or when they allow dynamics these are very limited to a particular kind of protein, and when they are flexible enough to allow all the above they are cumbersome to setup and execute and/or commercial products.

We believe HandMol is better than the available options, even in this version that we consider a prototype, because of everything it offers, its many modern features, its easy integration to existing programs, and its open use system. It is no surprise that, like all other apps and programs for molecular graphics and modeling in VR, HandMol is far from being a tool that can replace molecular graphics in research and possibly even in education. However, we do believe that HandMol might be useful enough to quickly assist users in carrying out the hardest, most 3D-intensive tasks, of which we have shown some examples. We discuss below specific potential applications in education and in chemistry/structural biology research. And we hear your ideas (and feedback and suggestions in general) at https://docs.google.com/forms/d/e/1FAIpQLSeC9rUEgJ_7eXhXqPE7r8FSg5qDvY_xGy-eTZSGqfBBuOYY4w/viewform?usp=sf_link.

### Web-based nature to maximize reach and accessibility

Central to our advocacy for web-based solutions as in MolecularWebXR,^5^ HandMol utilizes WebXR for VR molecular visualization and manipulation, in-browser speech recognition for oral commands and speech synthesis for auditory feedback, WebRTC for communication between instances, and calls to external APIs for molecular mechanics calculations. The modular integration of these technologies, and especially the WebXR API, allows HandMol to run on a variety of devices, including VR headsets, computers, tablets, and smartphones -the latter even producing immersive experiences as good as those of VR headsets if used with cardboard goggles.

Like with other web-based software, HandMol’s easy access, availability and cross-device compatibility extend the accessibility of molecular modeling and graphics to a broader audience. This inclusivity is essential for facilitating collaboration and engagement in both educational and research settings, especially in low-and mid-resource institutions.

### Modularity and flexibility

HandMol’s modularity is highlighted by its capacity to accommodate different types of calculations through APIs, with relatively little effort. The paper specifically demonstrates the integration of a Python version of ANI and of OpenMM for molecular mechanics calculations, showcasing the flexibility of the tool. OpenMM gives easy access not only to Amber forcefields as used here but also to other forcefields of the Charmm and Martini families, among others. Thus, it is rather straightforward to extend HandMol with more functionalities for molecular mechanics. Requiring some more work yet totally feasible, HandMol could be extended via APIs to simulate experimental observables in real time, that could for example be compared against experimental data as a user moves atoms and molecules.

### Example applications to educational and research settings

In education, HandMol can serve as a virtual kit for molecular modeling, allowing students to build molecules pretty much like when using tangible modeling kits, and also explore conformations, grasp 3D structures visually, investigate isomerism, simulate chemical reactions and interactions, etc. The tool’s versatility enables independent student work, collaborative sessions, or teacher-led demonstrations.

HandMol’s accessibility on various devices makes it suitable for different classroom setups. Students could work alone, each running HandMol-VR independently or in pairs by linking HandMol-VR and HandMol-Computer both running in VR headsets, or they could follow the explanations of a teacher who runs HandMol-VR in a headset, etc.

In research, HandMol could immediately finds applications in setting up systems for molecular simulations, building molecules with complex 3D features that are hard to draw in 2D, assisting in protein design as exemplified, testing conformational changes and functions depending on motions, and of course the simple visual inspections of molecular structures. The collaborative features of HandMol can be particularly valuable for research teams working on joint projects, enabling real-time interactions and discussions within VR environments far more engaging, immersive and interactive than what streaming the view of a regular molecular graphics program can allow.

## Conclusion

We open up HandMol to be tested for free, having showcased different ways to apply it to research and education. We hope HandMol can shape the way molecular modeling, chemistry and structural biology are conducted, taught and communicated; and inspire similar tools for other fields. Do please let us know in the survey forms whether and how HandMol worked for you and what features you would like to see implemented.

## Supporting information

Word version of main text

## Notes

https://go.epfl.ch/handmol

### Competing Interest Statement

The authors have declared no competing interest.

