## Supplementary material for "HandMol: Coupling WebXR, AI and HCI technologies for Immersive, Natural, Collaborative and Inclusive Molecular Modeling": Word version of main text

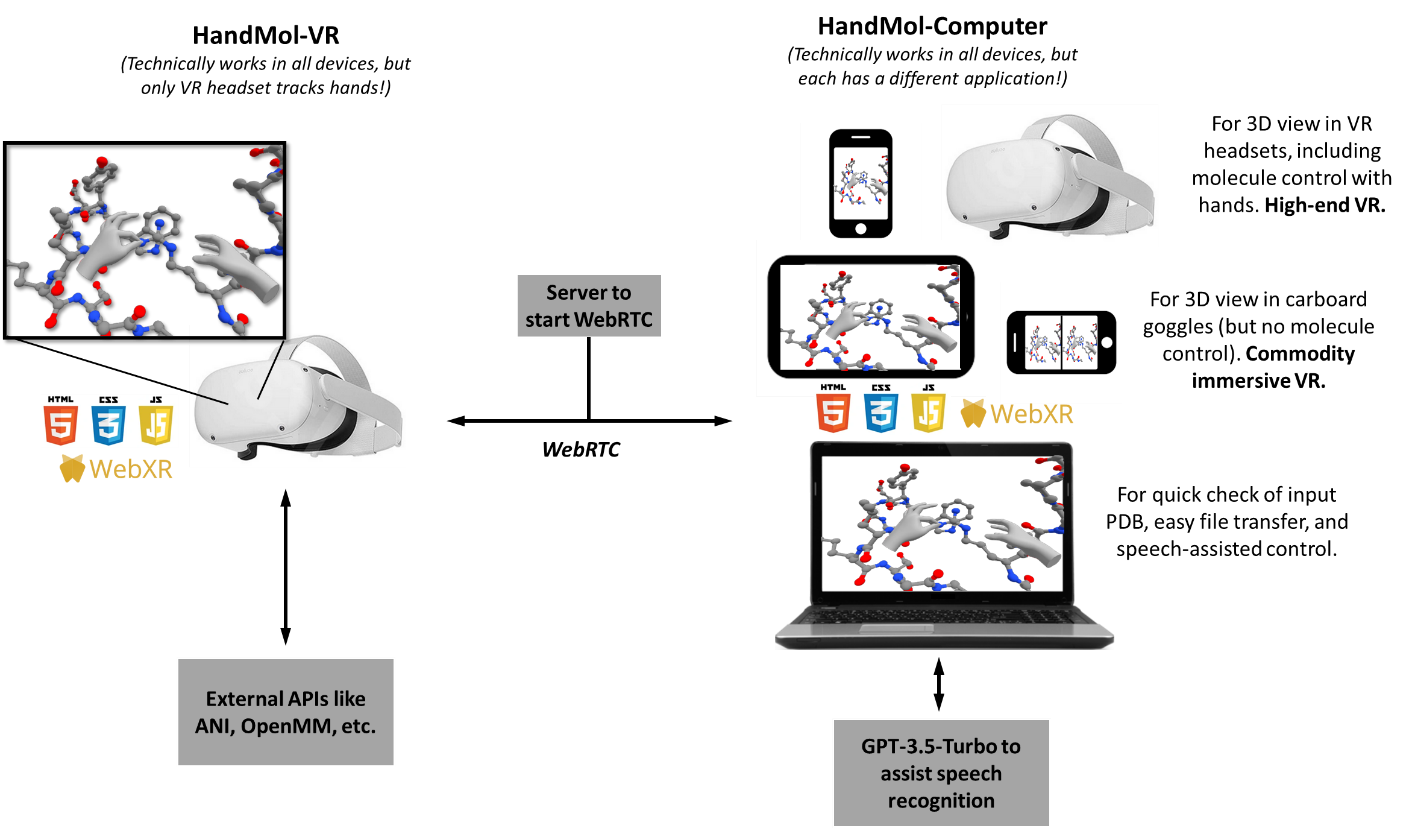

**Figure 1. HandMol architecture.** Scheme showing how HandMol’s components, called HandMol-VR and HandMol-Computer, are built from HTML, JS, CSS, WebXR, WebRTC, and API calling components; and how they can be used in different scenarios and with different pieces of hardware. The view is a true screenshot from inside a VR headset, where the background was removed and painted white for simpler visualization in this schematic figure. For the untouched screenshot, check Figure 2A; also check other figures of this article and try out the examples at <https://go.epfl.ch/handmol>.

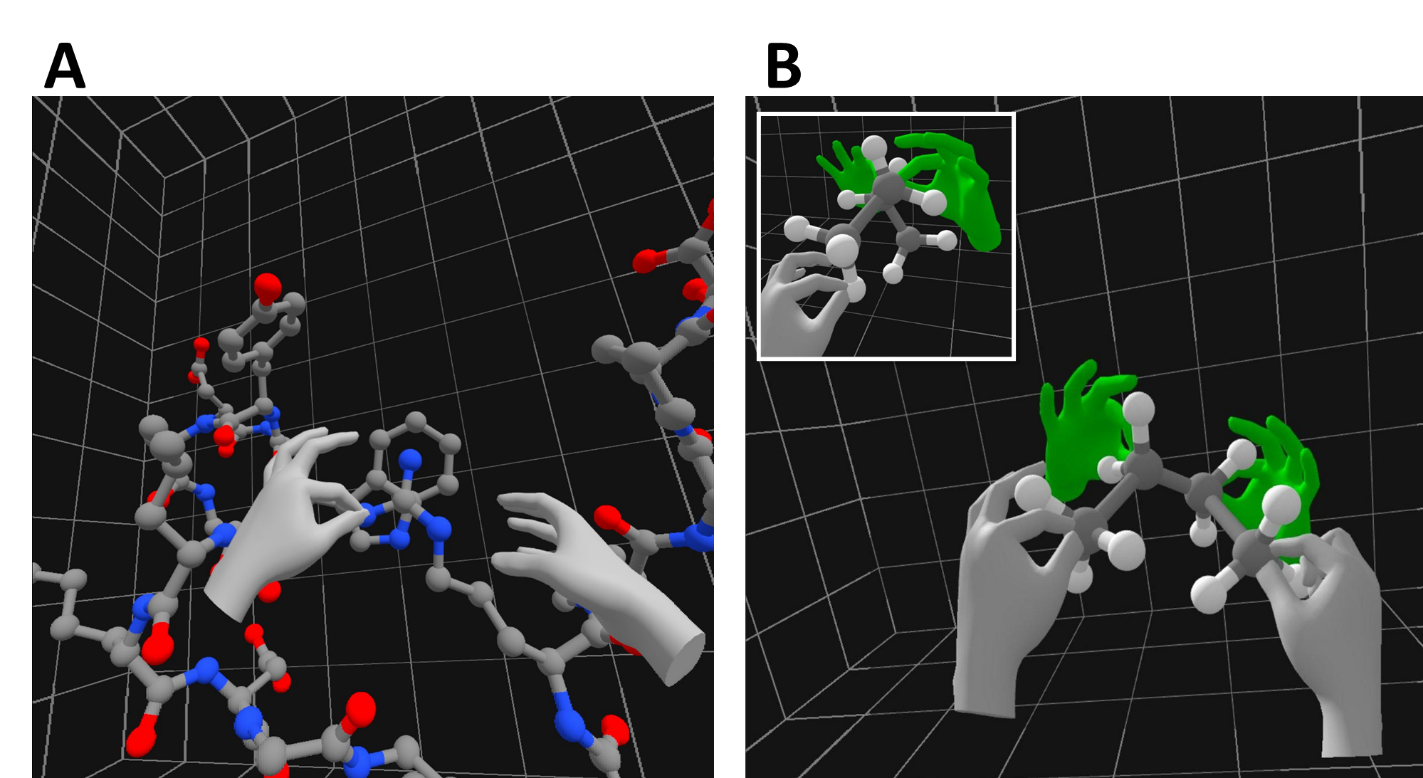

**Figure 2. One and two users inside VR with HandMol, and a chance to present more example systems.** (A) View of a user running HandMol-VR in a VR headset, here showing how an Arginine sidechain could form a cation-pi interaction with the aromatic rings of a Tryptophan sidechain. (B) View of a user running HandMol-VR in a VR headset and acting with his hands (seen in grey) on a molecular system together with another user who is running HandMol-Computer in a VR headset too (green hands). In this case, the users are together exploring the conformations of butane: while the green user holds two carbons tight, the grey user (whose view is shown) drives rotation around the central C-C bond. The inset shows the a similar operation but seen from a point aligned with the two central C atoms.

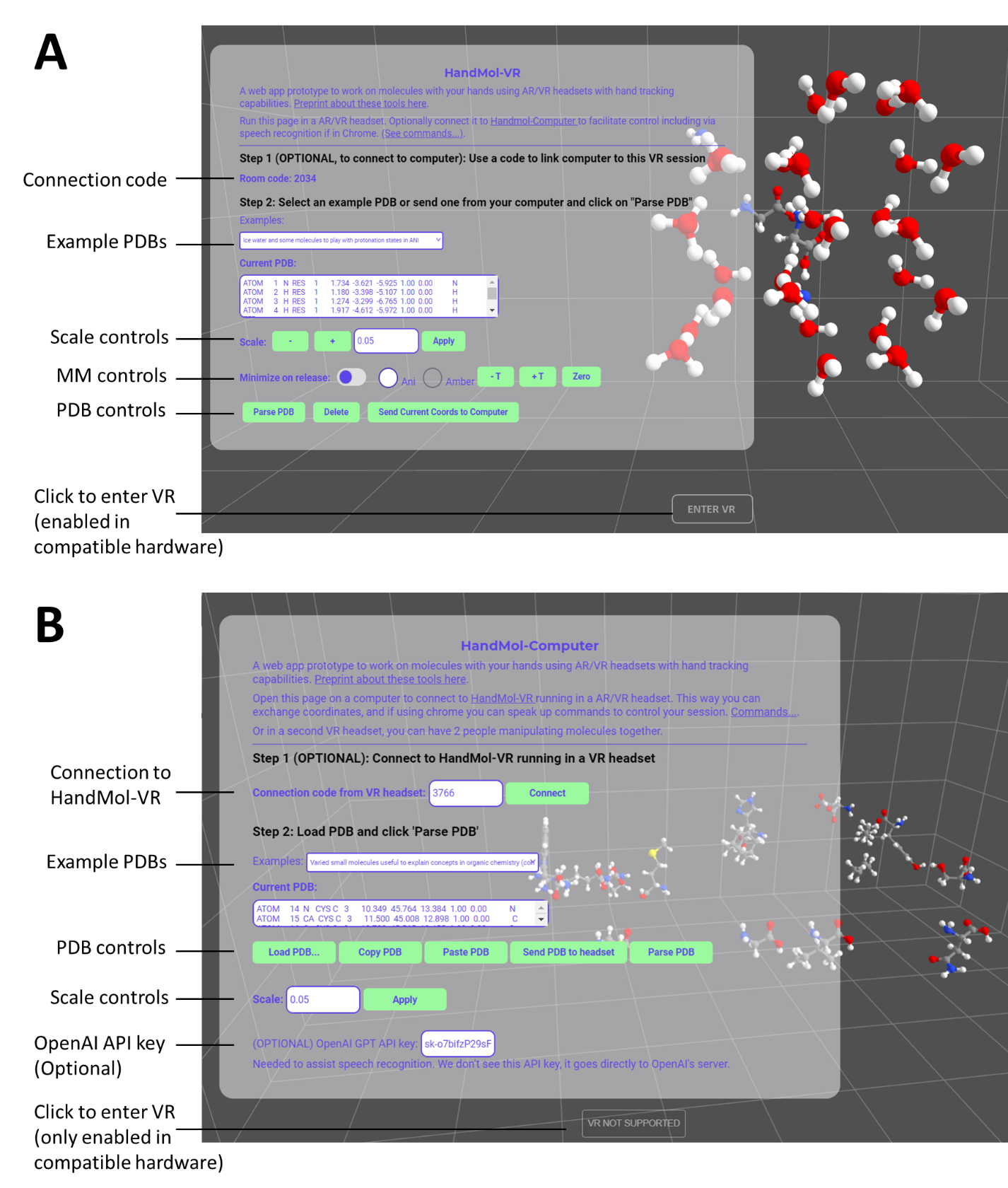

**Figure 3. HandMol-VR and HandMol-Computer outside VR mode, with controls exposed.** (A) HandMol-VR as seen in the web browser of a VR headset, before accessing VR (i.e. before clicking the *ENTER VR* button at the bottom). (B) HandMol-Computer as seen in the web browser of a laptop computer, which doesn’t allow VR access (hence the *ENTER VR* button displays as *VR NOT SUPPORTED*)*.*

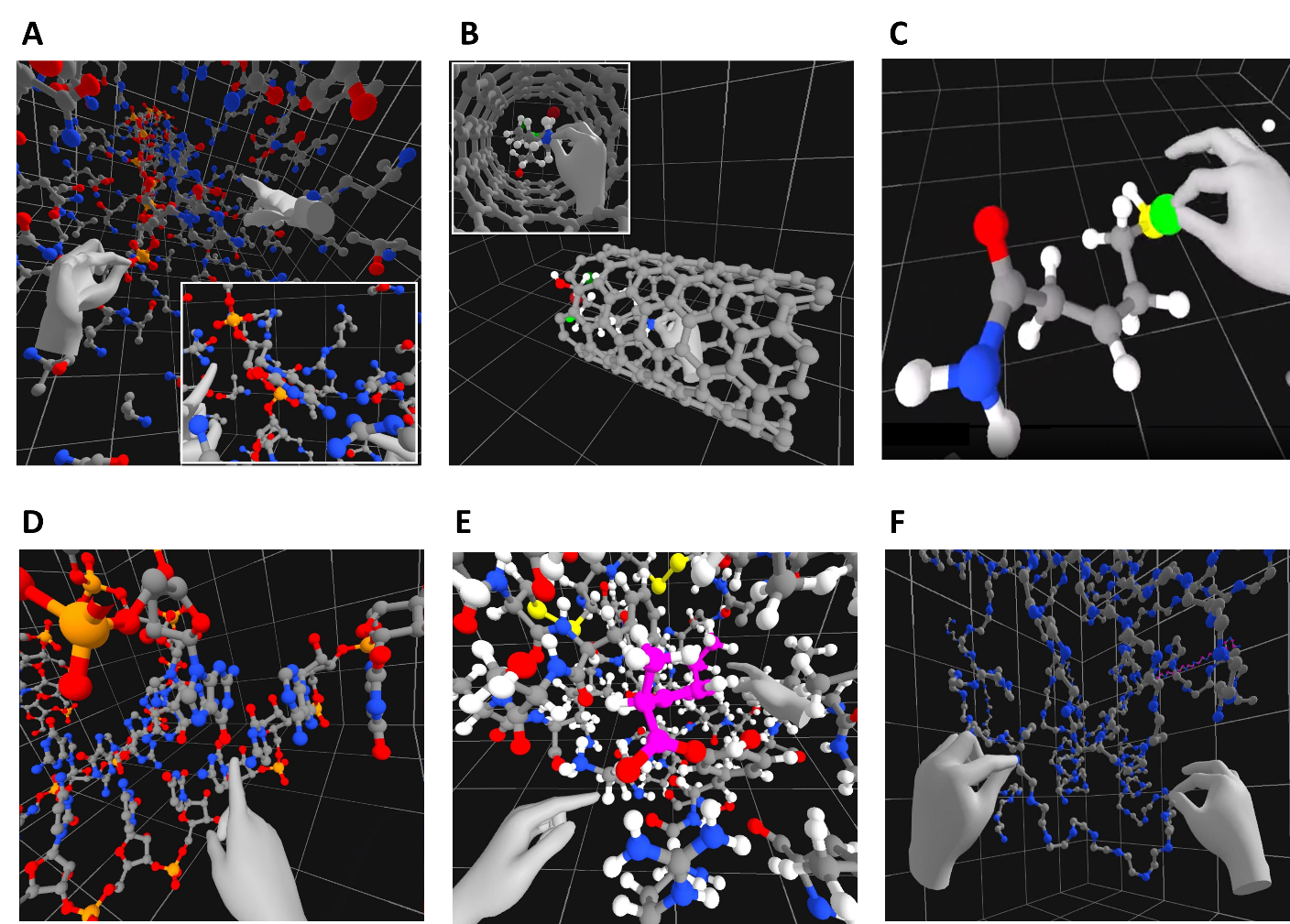

**Figure 4. Molecular mechanics in HandMol.** (A) A user threading a piece of single-stranded DNA through an aerolysin nanopore assisted by the rigid-body physics engine only, seen from the side where the DNA molecule is headed. The user’s left hand is seen pulling the DNA from a phosphate group, and the right hand is pointing at an Arginine that poised to interact with a DNA base at the reading site. Inset: View closer to the base being read, where the user’s right hand points at the Arginine’s guanidinium group and the left hand points at a phosphate that would be locked to other Arginine sidechains. (B) A carbon nanotube through which a user is pulling a molecule, assisted by the rigid-body physics engine. (C) Building a molecule inside the “Virtual Modeling Kit” example, here approaching an F atom to add it into the growing molecule, assisted by ANI-2x. See video version at <https://twitter.com/labriataphd/status/1726726066633343052>. (D) Pulling ibuprofen out of its pocket in BSA, assisted by rigid-body physics and running on a portion of an X-ray structure extracted around the small molecule. In this case the representation includes the H atoms, essential to better grasp how the small molecule fills the volume of the pocket and also helping to see the hydrogen bonds and electrostatic interactions the stabilize the complex. (E) Modeling how a peptide could connect two separated pieces of a protein being designed (the tool serving to approximate the length and pose of the required peptide, that will then be modeled in a program like RoseTTAFold Diffusion or similar). (F) A single-chain RNA molecule folded in space, explored with help of an AMBER forcefield.

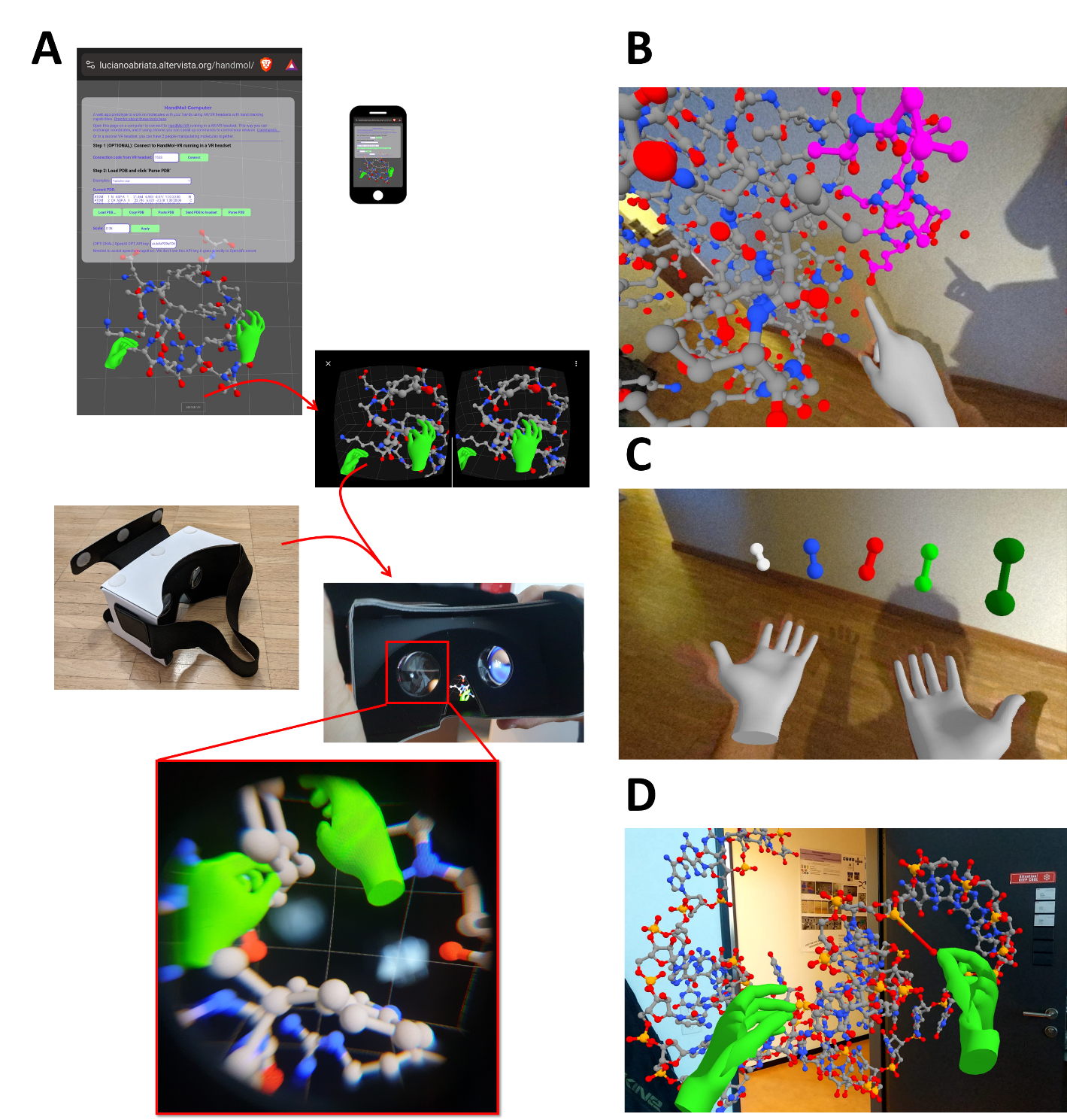

**Figure 5. Extended modes: VR in cardboard goggles, high-end mixed reality in modern VR headsets, and commodity augmented reality in smartphones.** (A) Accessing either HandMol-VR or HandMol-Computer in a smartphone (best if its browser is set in “Desktop site” to fit all the GUI in the screen) and clicking on *ENTER VR*, results in the WebXR API splitting the view in two for stereoscopy assisted by the device’s motion sensors. Inserting the smartphone inside cardboard goggles immediately allows immersive view much like in a VR headset, with the possibility to inspect the view by freely moving the head (but no displacement is possible, and no hand-tracking is available). (B) Visualizing a protein complex (Ubiquitin bound to an interacting helix, PDB 2D3G) with a VR headset (here Oculus Quest Pro) in mixed reality mode. (C) Building diatomic molecules made of different atoms assisted by ANI-2x, as seen in mixed reality. (D) HandMol-Computer running in Augmented Reality mode on a smartphone linked to another user who is running HandMol-VR in a headset, whose hands are seen in green.
